# Minimizing biosignal recording sites for hybrid noninvasive brain/neural robot control

**DOI:** 10.1101/2020.04.03.023325

**Authors:** Alessia Cavallo, Vincent Roth, David Haslacher, Marius Nann, Surjo R. Soekadar

**Affiliations:** Clinical Neurotechnology Laboratory, Neuroscience Research Center (NWFZ), Department of Psychiatry and Psychotherapy, Charité -University Medicine Berlin, Charitéplatz 1, 10117 Berlin, Germany; Applied Neurotechnology Laboratory, Department of Psychiatry and Psychotherapy, University Hospital of Tübingen, Calwerstr. 14, 72076 Tübingen, Germany

**Author notes:** Corresponding author: Surjo R. Soekadar.

**Keywords:** brain-computer interfaces, electroencephalography, electrooculography, rehabilitation robotics, robot control, wearable sensors

## Abstract

Noninvasive brain/neural controlled robots are promising tools to improve autonomy and quality of life in severe paralysis, but require biosignal recordings, such as electroencephalography (EEG) and electrooculography (EOG), from various sites distributed over the user’s head. This limits the applicability and practicality of noninvasive brain/neural robot control on an everyday basis. It would thus be very desirable to minimize the number of necessary recording sites paving the way for miniaturized, headset-like EEG/EOG systems that users with hemiplegia can mount by themselves. Here, we introduce a novel EEG/EOG brain/neural robot control strategy using only scalp electrodes placed near cortical sensorimotor areas. The strategy was tested across 16 healthy volunteers engaging in an EEG/EOG brain/neural control task. Classification accuracies were compared using scalp electrodes only vs. the conventional electrode placements across the scalp and face. To evaluate whether cranial muscle artifacts impede classification accuracy, participants were asked to chew during the task. We found that brain/neural classification accuracy was comparable and that chewing did not impact classification accuracies when using scalp electrodes only. Our results suggest that the proposed new strategy allows for reliable EEG/EOG-based brain/neural robot control, a critical prerequisite to broaden the use of noninvasive brain/neural assistive and rehabilitative technologies.

## I. Introduction

Introducing brain-controlled robotic devices from the lab into real-world applications is particularly challenging because environmental noise or other signal artifacts can impede classification of brain signals [1][2]. Recently, it was suggested that inclusion of additional physiological signals, e.g. related to eye or muscle movements, can improve safety and reliability of brain-controlled devices, such as active hand exoskeletons, when used outside the lab [3][4].

Besides providing an intuitive control mechanism for assistive applications, it was shown that translation of voluntary modulations of brain activity related to movement attempt or motor imagery into exoskeleton control can trigger neural recovery when used repeatedly [5][6][7]. This rehabilitative approach substantially extends the clinical relevance of brain-controlled devices, because approximately one in three stroke survivors suffers from severe motor impairments for which no other treatment strategy exists [8].

It would thus be very desirable to design brain/neural computer interaction (BNCI) systems that are broadly applicable and can be operated by stroke survivors without any assistance, e.g. in their home environment. This would not only increase the system’s impact on the user’s quality of life, but also foster generalization of learned skills into the user’s real-life environment.

A major obstacle towards this goal, however, relates to the necessary quantity of electrodes for recording biosignals from different recording sites across the user’s head. For instance, a well-established noninvasive brain/neural control paradigm uses electroencephalography (EEG) recorded from the scalp region as well as electrooculography (EOG) recorded from the medial or lateral canthus in close proximity to the eyes [9][10]. The lateral canthus is particularly suitable to record horizontal oculoversions (HOV), i.e. maximal horizonal eye movements, because of its proximity to the EOG’s electrical source (the corneo-retinal standing potential) and its distance to muscles of mastication (M. masseter, M. temporalis, and M. pterygoideus medialis et lateralis) that are broadly attached to the cranial bones. Electric field modulations related to eye movements recorded at the outer canthus typically range between 50-3500 μV [11] with their amplitudes inversely proportional to the square of the distance from the source [12]. Establishing a strategy that reduces the number and distribution of necessary electrodes, particularly of EOG electrodes placed into the user’s face, would pave the way for miniaturized, headset-like EEG/EOG systems that stroke survivors with hemiplegia can mount by themselves. Moreover, eliminating the necessity of electrodes in the facial region may improve user acceptance and therefore adoption and effectiveness. It was unclear, however, whether a reduction in recordings sites, e.g. for detection of HOV, is feasible without compromising brain/neural control performance.

Here, we investigated whether reliable EEG/EOG brain/neural robot control is feasible and robust using scalp electrodes placed in proximity to the sensorimotor areas only, and whether it provides similar brain/neural control performance as the conventional montage. While we reasoned that with increasing distance from the eyes classification accuracy of HOV would deteriorate and substantially worsen when muscles of mastication are contracted, we hypothesized that combination of several bilateral electrode pairs may compensate for such an effect and result in equivalent control performance.

## II. Materials and Methods

### A. Participants

16 BNCI-naive healthy volunteers (7 male, 9 females, mean age: 30.31 ± 9.24) were invited to the Department of Psychiatry and Psychotherapy at the Campus Charité Mitte (CCM) of the Charité – University Medicine Berlin, Germany, to participate in a 2-hour experimental session. All participants were righthanded [13], had no history of neurological or psychiatric disorders and did not take any medication on a regular basis. Before the experiment, all participants provided written informed consent. The study protocol was approved by the Charité’s local ethics committee (registration number EA1/077/1).

### B. Biosignal recordings

A custom version of the BCI2000 software platform *(www.bci2000.org)* was used for signal processing and visual feedback presentation on a computer screen. An active wet electrode system (actiCAP^®^) coupled with an amplifier unit (LiveAmp^®^, Brain Products GmbH, Gilching, Germany) was used to record and amplify biosignals from the following sites according to the international 10/20 system: F3, F4, T3, T4, C3, C4, P3, P4, and Cz (scalp electrodes). Moreover, in accordance to the standard EOG placements, two electrodes were placed at the left and right outer canthus (facial electrodes). To detect activity of muscles of mastication, two additional electrodes were placed on the right masseter muscle (Fig. 1). A reference electrode was placed at FCz and a ground electrode was placed at FPz. All biosignals were sampled at 1000 Hz and high-pass filtered at 0.1 Hz with a notch-filter at 50 Hz.

**Fig. 1.**
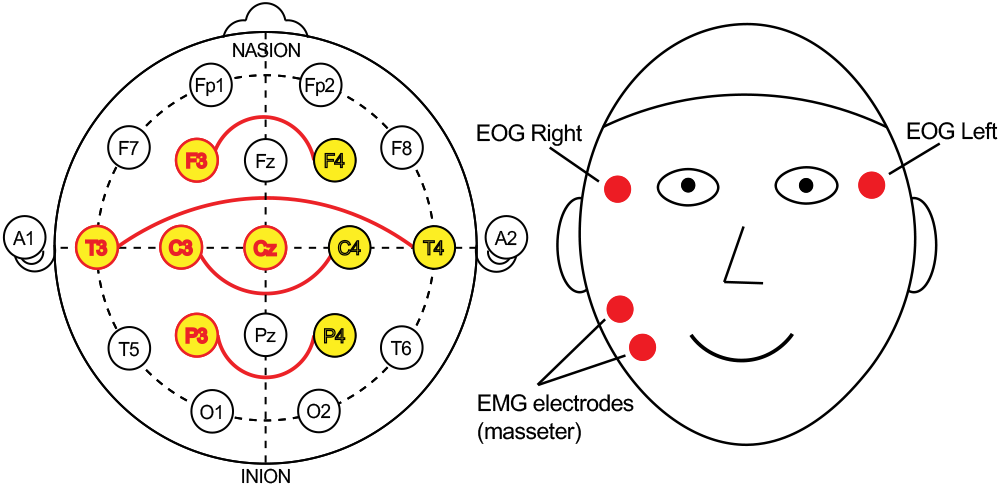
Distribution of biosignal recording sites across the scalp (left panel) and face (right panel). While signals from electrode sites covering the scalp and face were used in condition1, HOV detection was performed using scalp electrodes only in condition2 (indicated in yellow). To assess activity of muscles of mastication, two additional electrodes were placed over the right M. masseter.

#### 1) Signal processing chain for brain activity (SP_Brain_)

For online processing of brain activity, the signals from F3, T3, C3, P3 and Cz were filtered between 1 and 30 Hz using a second order Butterworth filter (Fig. 1). A surface Laplacian filter was applied according to [14]. The power spectrum of a 0.4 s window was computed at 10-12 Hz using an autoregressive model of order 100 with 15 evaluations per bin. The frequency range was chosen to detect event-related synchronization/ desynchronization (ERS/ERD) of sensorimotor rhythm (SMR) typically occurring during motor imagery or motor planning. The output of this processing chain (SP_Brain_) was translated into online visual feedback (see section II.C). Before BNCI online control, a calibration run was performed to compute an individualized SMR-ERD detection threshold for each participant.

##### a) Calibration of the SMR-ERD detection threshold

Participants were asked to imagine to close their right hand when the word “close” was displayed on a computer screen placed in front of them. When the word “relax” was displayed, participants were instructed to rest. “Close” and “Relax” visual cues were presented 10 times each with a duration of 5 s followed by an inter-trial interval (ITI) of 4 s. The average output of the SP_Brain_ during all cues was calculated and used as a reference value (RV). To obtain the relative deviation from the RV during each trial, RV was subtracted from the chain output and the result divided by RV. The SMR-ERD detection threshold was then set to the average of the chain output during ten “close” calibration trials and kept unchanged throughout the experiment.

#### 2) Signal processing chain for HOV (SP_HOV_)

To detect HOV, four bipolarized electrode pairs were used (F3-F4, T3-T4, C3-C4, P3-P4), subtracting signals from electrodes on the right hemisphere form signal from electrodes on the left hemisphere (Fig. 1). The bipolar signals were then low-pass filtered at 1.5 Hz using a second order Butterworth filter and summed together in order to increase the signal-to-noise ratio. The output of SP_HOV_ was translated into visual feedback (see section II.C). Before BNCI online control, a calibration run was performed to calculate the HOV detection threshold for each participant.

##### a) Calibration of the HOV detection threshold

Participants were instructed to perform maximal HOV without turning their head when an arrow to the left or the right was displayed on the computer screen. Each arrow was presented five times for a duration of 2 s and was followed by an ITI of 4 s. For each arrow to the left (r. right), the maximum (r. minimum) of the SP_HOV_ was computed. The detection threshold for left (r. right) HOV was set to two third of the average maximum (r. minimum) chain output and was kept unchanged throughout the experiment.

The calibration runs and subsequent calculation of the SMR-ERD detection and HOV thresholds were repeated when BNCI control remained below chance level at the end of calibration.

### C. Testing the BNCI system

The experimental design included two blocks (block1, block2) with two runs each (run1, run2) comprising a total of 160 trials. While in block1, participants were instructed to relax their cranial muscles, participants were asked to activate muscles of mastication by clenching their teeth in block2. During each run, a sequence of 40 trials was presented. To allow for calculation of sensitivity and specificity of BNCI control, four different visual cues were displayed ten times in a randomized order. Each cue was displayed for 5 s followed by an ITI of 4s (Fig. 4). Before the start of the experiment, participants were familiarized with the experimental design and procedure.

1. During the first cue (cue1, Fig. 4), a yellow half circle was displayed in conjunction with the word “close” to indicate that the participant should imagine right-hand closing motions. If the output of SP_Brain_ fell below the SMR-ERD detection threshold, the yellow circle filled by an additional 6%. The yellow circle was fully filled when the SMR-ERD detection threshold was exceeded for more than 3 s.
2. During the second cue (cue2, Fig. 4), a yellow circle accompanied with the text “open” was displayed to indicate that the participant should execute an HOV. If the SP_HOV_ exceeded (r. undercut) the HOV detection threshold, the yellow circle changed into a yellow half circle.
3. During the third cue (cue3, Fig. 4), a blue bar was displayed accompanied by the word “relax” to indicate that the participant should rest. If output of the SP was below the SMR-ERD detection threshold, the blue bar increased in size, while decreased in size when the output exceeded the threshold.
4. During the fourth cue (cue4, Fig. 4), a yellow circle accompanied by the word “relax” was displayed to indicate that the participant should rest without HOV. If the output of SP_HOV_ exceeded (r. undercut) the detection threshold for left (r. right) HOV, the yellow circle changed to a yellow half circle.

**Fig. 2.**
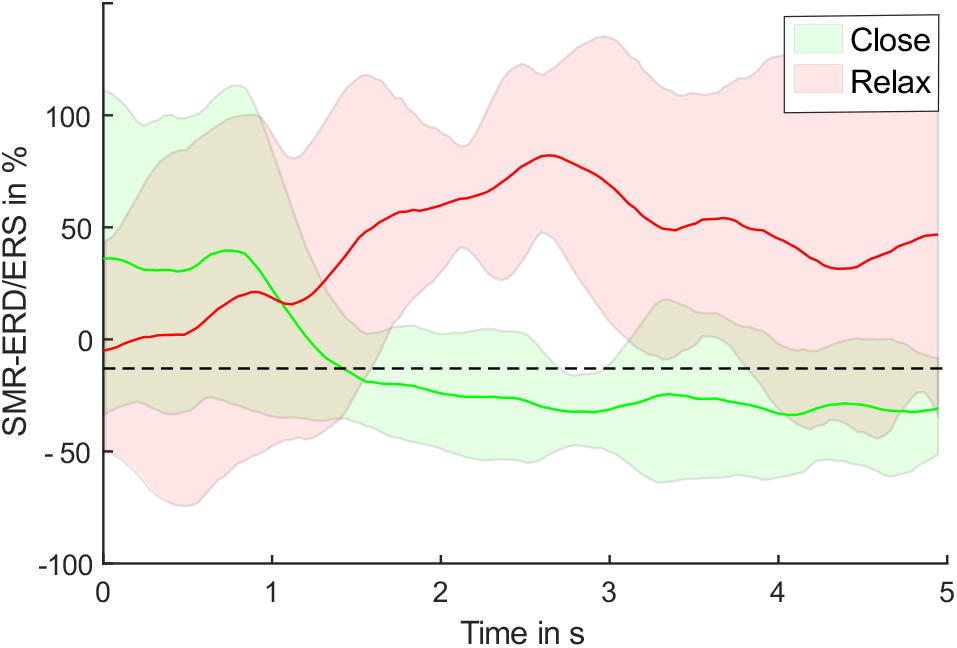
Illustration of sensorimotor rhythm event-related desynchronization and synchronization (SMR-ERD/ERS) in a representative participant. The red line represents SMR-ERS during the instruction to relax, while the green line represents SMR-ERD during the instruction to imagine hand closing motions. The SMR-ERD detection threshold is indicated as black dotted line.

**Fig. 3.**
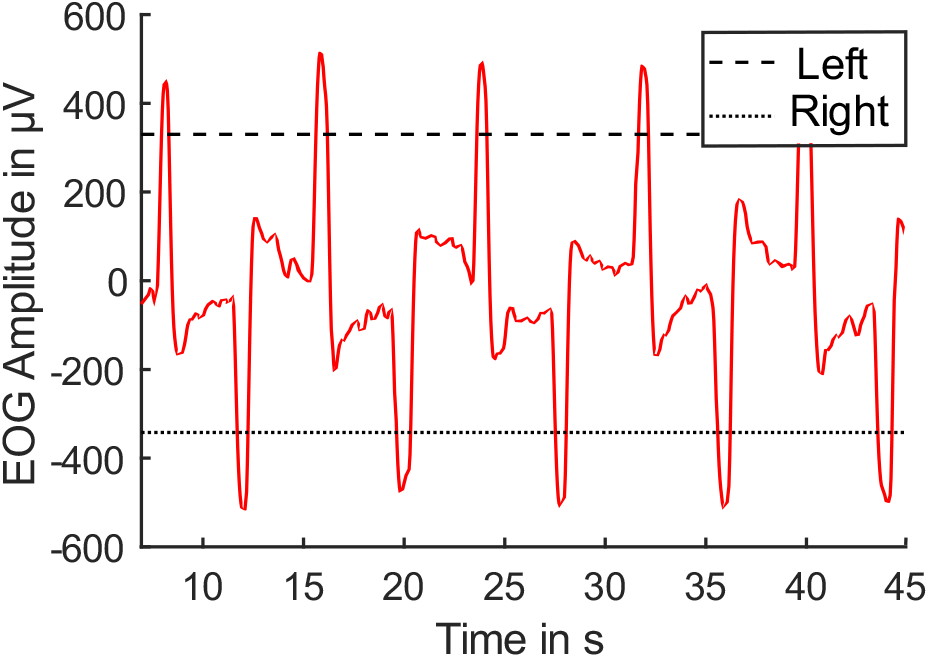
Illustration of horizontal oculoversions (HOV) to the left (positive deflections) and right (negative deflections) as measured by scalp electrodes only (condition2) in a representative participant. The HOV detection threshold for BNCI control is indicated as a black dashed line for HOV to the left, and a dotted line for HOV to the right.

**Fig. 4.**
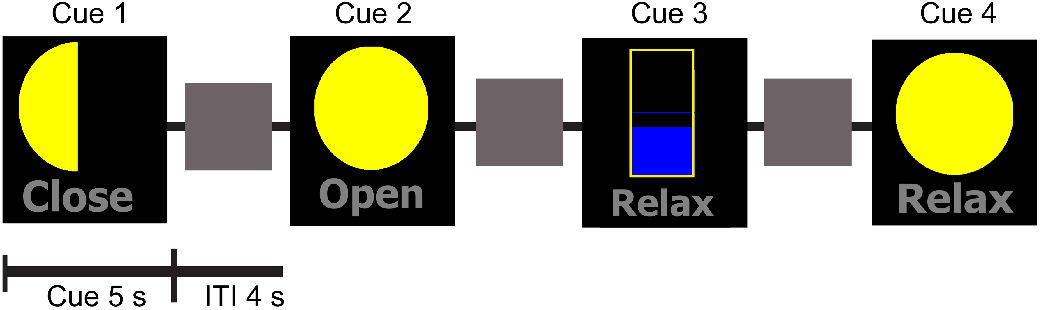
Task instructions were provided using four different graphical cues. Cue1 (left panel) indicated that the participant should engage in motor imagery, cue2 indicated that the participant should perform a horizontal oculoversions (HOV). Cue3 and cue4 indicated that participant should rest and relax.

### D. Offline data analysis

The overall BNCI control performance for each participant and block was calculated as the average of the following four performance values:

1. The BNCI system’s sensitivity (true positive rate) for SMR-ERD detection was computed as the mean percentage of yellow circle fillings achieved during 20presentations of cue1 (“close” instruction accompanied by showing a yellow half circle).
2. The BNCI system’s sensitivity for HOV detection was computed as the percentage of circles that changed from a full to a half yellow circle during 20 presentations of cue2 (“open” instruction accompanied by showing a yellow full circle).
3. The BNCI system’s specificity (true negative rate) for SMR-ERD detection was computed as the mean percentage of time the blue bar decreased in size during 20 presentations of cue3 (“relax” instruction accompanied by showing a blue bar).
4. The BNCI system’s specificity for HOV detection was computed as the percentage of trials in which yellow full circles that did not switch to yellow half circles during 20 presentations of cue4 (“relax” instruction accompanied by showing a yellow full circle).

To ensure that during block2 only trials were included in which participants activated their muscles of mastication, only trials were included in which EMG signals (10-100 Hz) that were recorded over the right masseter exceeded two standard deviations of EMG signals recorded during block1. To calculate the overall control performance when using the conventional electrode montage, i.e. scalp and face electrodes vs. scalp electrodes alone, BNCI control performance was calculated offline for both conditions (condition1: scalp & face, condition2: scalp only). In analogy to SP_HOV_ described in II.C.2, corresponding electrodes were bipolarized and signals filtered from 0.1 to 1.5 Hz using a second order Butterworth filter. The threshold for left (r. right) HOV detection was then calculated using the calibration run. The BNCI system’s sensitivity and specificity for HOV detection in condition1 and condition2 as well as the overall BNCI system’s control performance was then computed offline.

To test whether the overall control performance in condition1 was inferior to condition2, a one sample-paired t-test was performed for block1 and block2. Non-inferiority was assumed at a difference smaller than 2 % with p < 0.05.

## III. RESULTS

After familiarization with the task, all participants were able to voluntarily generate SMR-ERD and HOV for brain/neural control. In block2, in average 2.01 ± 1.7 % of cue2 and cue4 were discarded due to insufficient masseter activity during the trials.

In block 1, there was no difference in overall BNCI control performance between condition1 and condition2 (p < 0.001) (Fig. 6). Overall BNCI control performance in condition1 was 87.68 ± 31.48 % and 87.91 ± 31.88 % in condition2 indicating that EEG/EOG detection was comparable when using scalp electrodes only as compared to the conventional electrode montage including scalp and face areas.

**Fig. 5.**
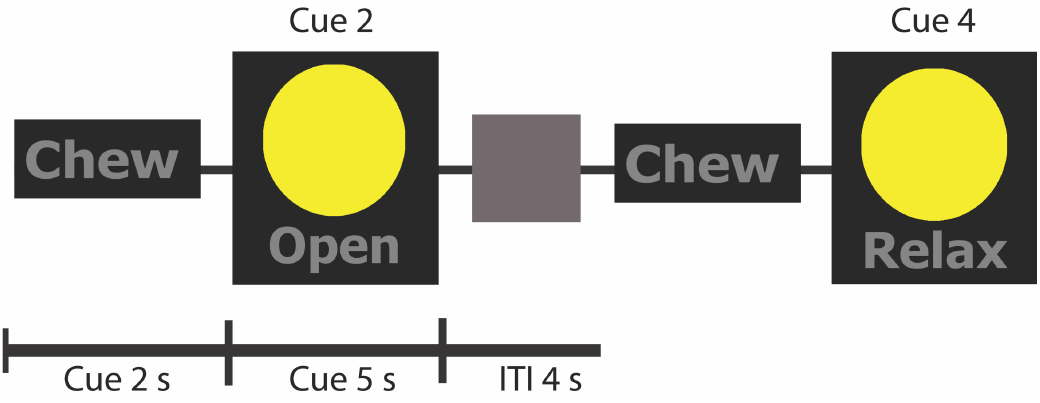
In block2, an additional visual cue (chew) was used to indicate that the participant should clench the teeth to activate muscles of mastication

**Fig. 6.**
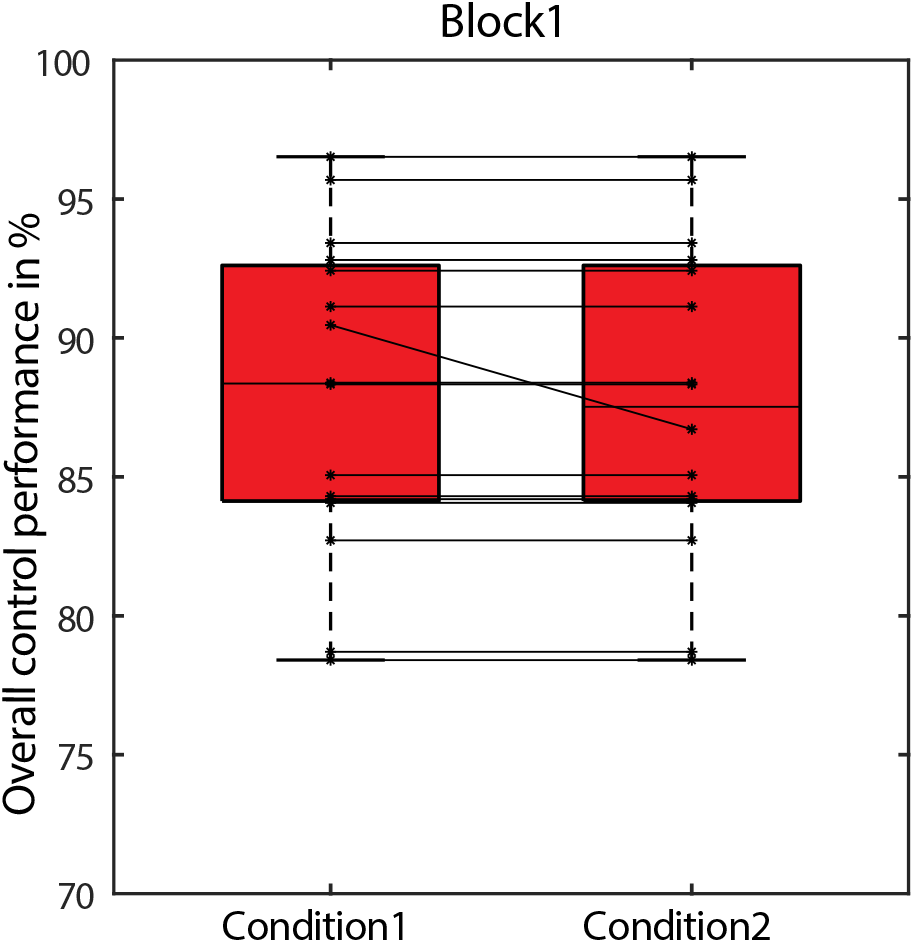
Overall BNCI control performance across all study participants in block1 under condition1 (using biosignals from electrode sites distributed over the scalp and face) (left) and condition2 (using biosignals from the scalp region only) (right). Changes in BNCI performance between conditions is shown as black line for each participant. The median is shown as block line in each box graph, the 25 th and 75 th percentiles are indicated by the error bars.

There was no difference in BNCI control performance in block2 in which participants engaged in mastication (p < 0.001) (Fig. 7). Overall BNCI control performance reached 87.17 ± 32.52% with scalp & facial electrodes (condition1) and 86.74 ± 39.12% with scalp electrodes only (condition2) (Fig. 7).

**Fig. 7.**
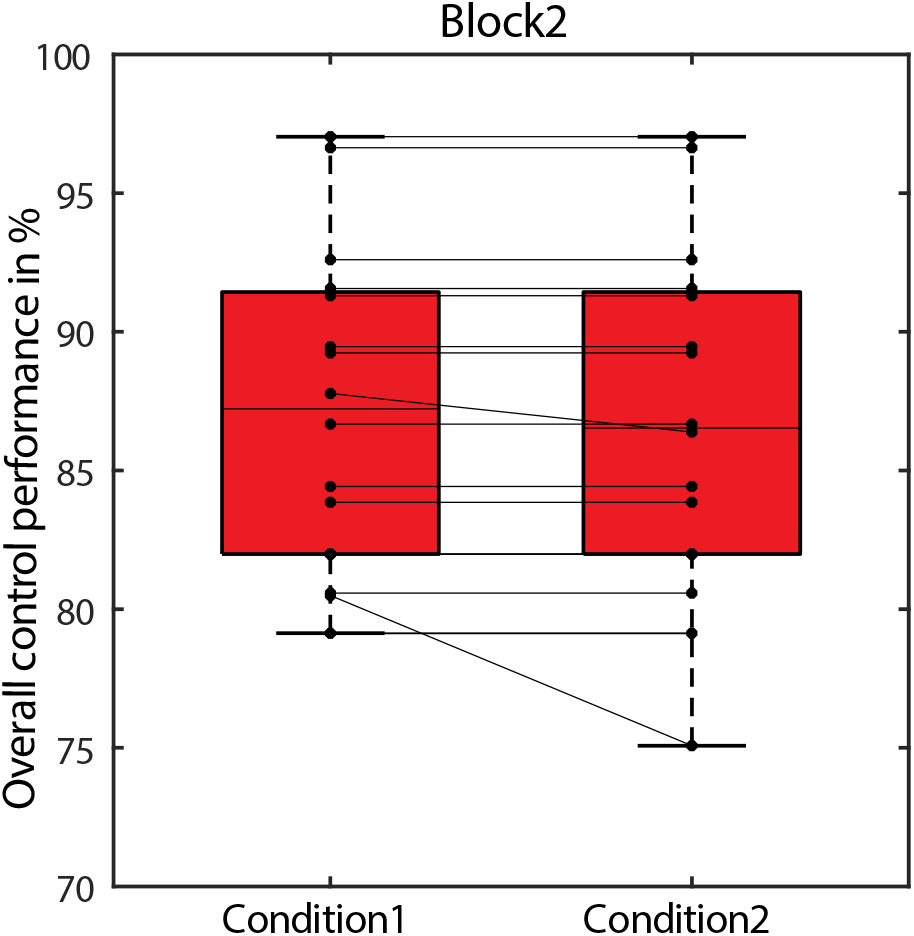
Overall BNCI control performance across all study participants in block2 under condition1 (using biosignals from electrode sites distributed over the scalp and face) (left) and condition2 (using biosignals from the scalp region only) (right). Changes in BNCI performance between conditions is shown as black line for each participant. The median is shown as block line in each box graph, the 25 th and 75 th percentiles are indicated by the error bars.

Under condition1, sensitivity and specificity of HOV detection across all participants and blocks reached 100%. Under condition2, 100% sensitivity and specificity of HOV detection was reached by 15 participants in block1 and 14 participants in block2. While specificity of HOV detection was 100% across all participants, sensitivity was 85% in block 1 of one participant, and 95% and 78% respectively in block 2 (see Fig. 7).

## IV. DISCUSSION

Reducing the number of biosignal recording sites for noninvasive hybrid brain/neural robot control is important to broaden applicability of assistive and rehabilitative BNCI systems. Here, we tested a strategy for reliable EEG/EOG brain/neural control that exploits the fact that the eye’s electric fields spread over the scalp and can be detected over the brain’s sensorimotor areas [12]. It was unclear, however, whether online HOV detection can be achieved at similar accuracies using this strategy compared to the established approach recording HOV from the outer canthus [4][9][10][15]. We found that EEG/EOG classification accuracies were comparable when using scalp electrodes placed in proximity to the sensorimotor areas only (condition2) as compared to the conventional approach with additional electrodes placed in the facial area at the outer canthus (condition1). Robustness of BNCI classification in each condition was evaluated by comparing classification accuracies with study participants relaxing their jaw (block1) vs. activating their muscles of mastication by clenching their teeth (block2). Independent of masseter contractions, classification accuracies were comparable across blocks indicating that the new strategy is robust even when engaging in mastication involved in various activities of daily living, such as eating, drinking or speaking. While specificity of HOV detection was 100% across all participants, blocks and conditions, sensitivity was 100% for 15 participants under condition2 in block1 and 14 participants in block2. The reason for this reduction in sensitivity in block2 found in two participants was most likely due to a rather conservative adjustment of the HOV detection threshold. Given that specificity was 100% across all participants, it is conceivable that setting the HOV detection threshold not at two third of the average maximum or minimum of SP_HOV_, but slightly below that (e.g. at 60%), would result in 100% sensitivity across all participants without affecting specificity of HOV detection. This issue should be further investigated in follow-up studies.

While this study focused on reducing the number of head areas required for reliable BNCI control (scalp & face vs. scalp only), it is conceivable that further reduction of the number and distribution of electrode recording sites is possible. Here, use of tri-polar concentric ring electrodes could be a viable solution [16][17]. It was suggested that individualization of EEG electrode sites may improve classification accuracies [9], particularly in patients who suffered a stroke [18] or other conditions associated with cortical reorganization [19]. In this context, it will be important to assess the impact of such individualization on EEG/EOG classification accuracies when using the proposed strategy. Also, it is unclear how different disease conditions that are associated with eye movement disorders, e.g. Parkinson’s or multiple sclerosis [20], affect reliability of the proposed approach.

Besides minimizing the number of recording sites, also other factors will influence applicability and user acceptance of BNCI systems. Here, wearing comfort and a non-stigmatizing overall design are key elements to increase adoption of BNCI technology into everyday life environments. Future assistive and rehabilitative BNCI systems should be as intelligent, but also as intuitive and simple as possible [21]. Moreover, various neuroethical dimensions have to be considered when bringing BNCI technology out of the lab into real world applications [22].

Besides improving neuroliteracy, ensuring accountability, responsibility, privacy and security are important dimensions on this path. The proposed BNCI control strategy specifically contributes to the dimension of accountability, because it provides a reliable option to, e.g., stop (veto) an unwanted action of a brain-controlled actuator. Such veto function is particularly essential for shared control paradigms, in which a human interacts with an autonomous robotic system [23]. The proposed veto function could be easily implemented in existing EEG-based BCI paradigms without the necessity of adding EOG electrodes in the facial region and thus increase their applicability in real-world scenarios. Further validation of this approach in clinical studies involving patient populations will be necessary, however.

